# Temporal dynamics of energy-efficient coding in mouse primary visual cortex

**DOI:** 10.1101/2024.12.17.628997

**Authors:** S. Amin Moosavi, Antonia Pastor, Alfredo G. Ornelas, Elaine Tring, Dario L. Ringach

**Affiliations:** Departments of Neurobiology, David Geffen School of Medicine, University of California, Los Angeles, Los Angeles, CA 90095; Departments of Psychology, David Geffen School of Medicine, University of California, Los Angeles, Los Angeles, CA 90095; Interdepartmental Neuroscience Program, Brain Research Institute, David Geffen School of Medicine, University of California, Los Angeles, Los Angeles, CA 90095

## Abstract

Sparse coding enables cortical populations to represent sensory inputs efficiently, yet its temporal dynamics remain poorly understood. Consistent with theoretical predictions, we show that stimulus onset triggers broad cortical activation, initially reducing sparseness and increasing mutual information. Subsequently, competitive interactions sustain mutual information as activity declines and sparseness increases. Notably, coding efficiency, defined as the ratio of mutual information to metabolic cost, progressively increases, demonstrating the dynamic optimization of sensory representations.

---

Efficient coding posits that sensory systems optimize the representation of natural stimuli by reducing redundancy while preserving information^1–4^. Simulations have shown that optimizing the sparse representations of natural images in a linear model leads to receptive fields similar to those found in the primary visual cortex (V1)^5^. These findings established efficient coding as a compelling, normative theory for understanding cortical function, prompting rigorous analyses of its properties^6–10^.

Subsequent experimental studies offered additional support for this framework. Prior work has shown that individual V1 responses are sparse in response to natural images^11^ and that sparseness increases when V1 is stimulated with patches of increasing size^12^, suggesting that sparsification relies on contextual information^13,14^. Moreover, population responses are sparser when V1 is stimulated with natural scenes as compared to synthetic stimuli, enhancing their discriminability^15^. Other theoretical proposals, such as the normalization model^14^, also reduce redundancy to produce more efficient responses^3,13^. Mechanistic models, such as soft winner-take-all networks, have been proposed to explain how a neural network may dynamically refine its activity, progressively increasing sparseness during sensory responses^16–18^.

To study the temporal evolution of efficient representations in cortical populations, we used two-photon imaging to measure the response of V1 neurons to visual stimulation (**Fig 1a**) (**Methods**). In the first experiment, the stimulus consisted of a sequence of flashed, sinusoidal gratings with orientations uniformly sampled between 0 and 170 deg in 10 deg increments, yielding 18 possible orientations (**Fig 1b**). Each grating was flashed for 1.5 sec with no gaps between stimuli. In the second experiment, the stimulus consisted of a sequence of 1 sec, natural movie clips, randomly selected from a set of 18 distinct segments, once again, with no gaps between them (**Fig 1c**). For both experiments, the 18 orientations or movie clips were treated as distinct stimulus “classes”.

**Figure 1.**
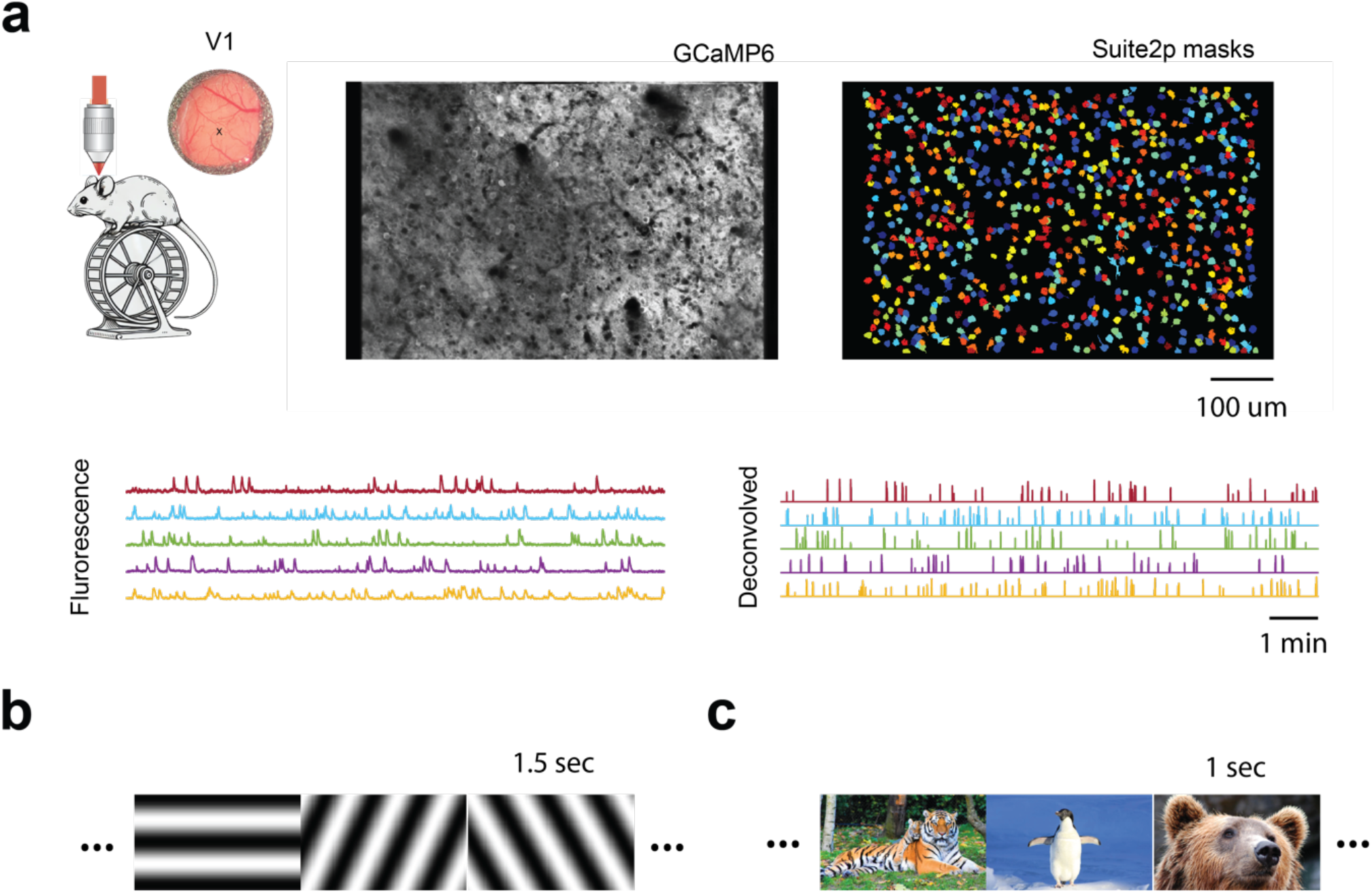
Experimental setup and visual stimulus. **a**. Two photon imaging was used to measure the activity of neurons in primary visual cortex. Images were registered, cells segmented, raw fluorescence extracted and deconvolved using Suite2p28. **b, c**. The visual stimulus consisted of a sequence of sinusoidal gratings or nature movie clips. The former was presented at a rate of one per 1.5 sec, while movie clips lasted for 1 sec. Stimuli were presented back-to-back with no blanks in-between.

We aimed to determine whether the temporal dynamics of sparse coding align with the adaptive optimization of energy-efficient representations. A sparsification network predicts that, at the onset of a new stimulus in a sequence, population activity will rise rapidly due to broad network activation, starting from a state shaped by the late responses to the preceding stimulus. This initial activity, driven by neurons broadly tuned to various features of the new stimulus, is expected to result in a transient decrease in sparseness. Concurrently, the ability to classify the stimulus, measured by mutual information, should increase as the signal-to-noise ratio of the response improves with the rise in activity. In the late phase of the response, competitive interactions within the population are expected to suppress redundant activity, increasing sparseness while maintaining high mutual information and thereby optimizing representation efficiency. This study aimed to test whether these predicted relationships are observed in population responses in mouse V1.

To analyze the data we derived three key metrics: metabolic cost, population sparseness^19^, and mutual information (**Methods**). Metabolic cost, *μ*(*t*), was defined as the total population activity at time *t* after stimulus onset^20^. Population sparseness, *s*(*t*), was quantified using the Gini index^21^, which ranges from zero (when all the neurons are firing equally) to one (when all but one neuron is active). As an alternative index of sparseness, we also used the percentage of active neurons, *a*(*t*). Finally, adjusted mutual information^22^, *I*(*t*), was estimated based on cross-validated decoders trained at different times after stimulus onset^15^, which ranges from zero (where there is no association between stimulus and response) to one (for perfect association).

Results from individual experiments using sinusoidal gratings reveal consistent patterns in the relationship between these metrics (**Fig. 2**). The dynamics exhibit two distinct phases. In the early phase, stimulus onset triggers a rapid increase in metabolic cost, which peaks shortly thereafter, consistent with prior studies^23^. We define the time-to-peak as the moment when this maximum value is reached, and the early phase of the response as the period from stimulus onset to this time (**Fig 1**, blue shaded rectangle). During this phase, sparseness decreases, while mutual information increases. In the late phase, which follows the time-to-peak, metabolic cost declines to baseline levels, sparseness returns to baseline after a slight overshoot, and mutual information decreases modestly but remains consistently high throughout the response. The percentage of active neurons at any time is highly correlated with the sparseness measure as defined by the Gini index.

**Figure 2.**
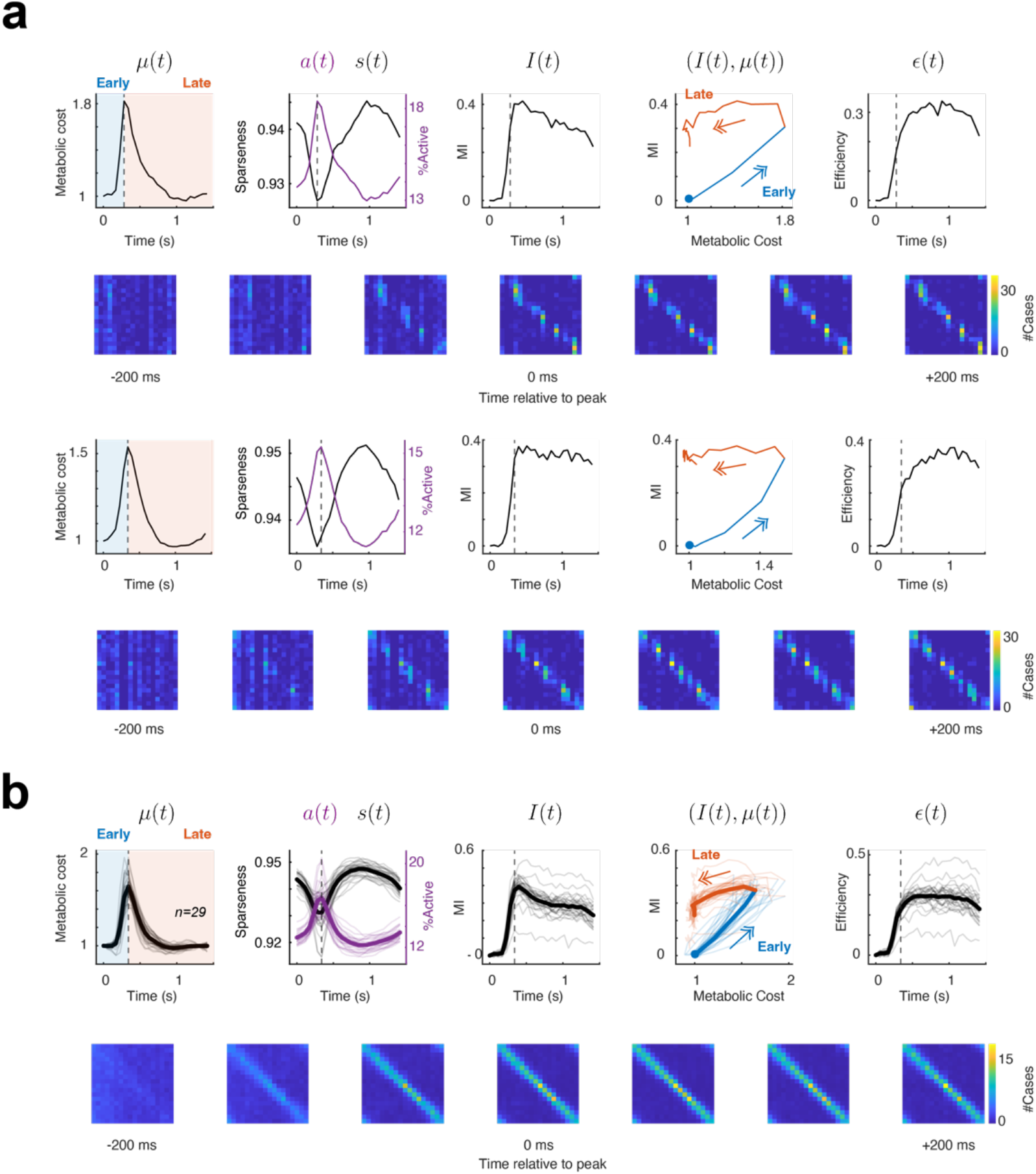
Dynamics of efficient coding using synthetic stimuli. **a**. Results in two individual sessions. The panels illustrate, from left to right: metabolic cost, *μ*(*t*); sparseness and fraction of activity neurons, *s*(*t*) and *a*(*t*) respectively; mutual information, *I*(*t*); the trajectory in the (mutual information, metabolic cost) plane; and efficiency, *∈*(*t*). In the metabolic cost panel, shaded rectangles show the early (blue) and late (red) phase of the response, delineated by the time-to-peak (vertical dashed line). The point of the trajectory at time zero is indicated by a blue dot, with arrows indicating the direction of the trajectory. The trajectory segment in blue shows the path during the early phase of the response, while the segment in red shows the one in the late phase. Arrows show the direction of time, with the blue dot denoting the onset of stimulation. Images show the confusion matrix at different times relative to the time to peak at 0 ms. Rows represents the true class of the stimulus, and the columns represent the predicted class. **b**. Population data across all the experiments (*n=19* sessions). Transparent curves show the result of individual experiments while the solid curves show the average response across sessions. The images show the average confusion matrices relative to the time to peak.

**Figure 3.**
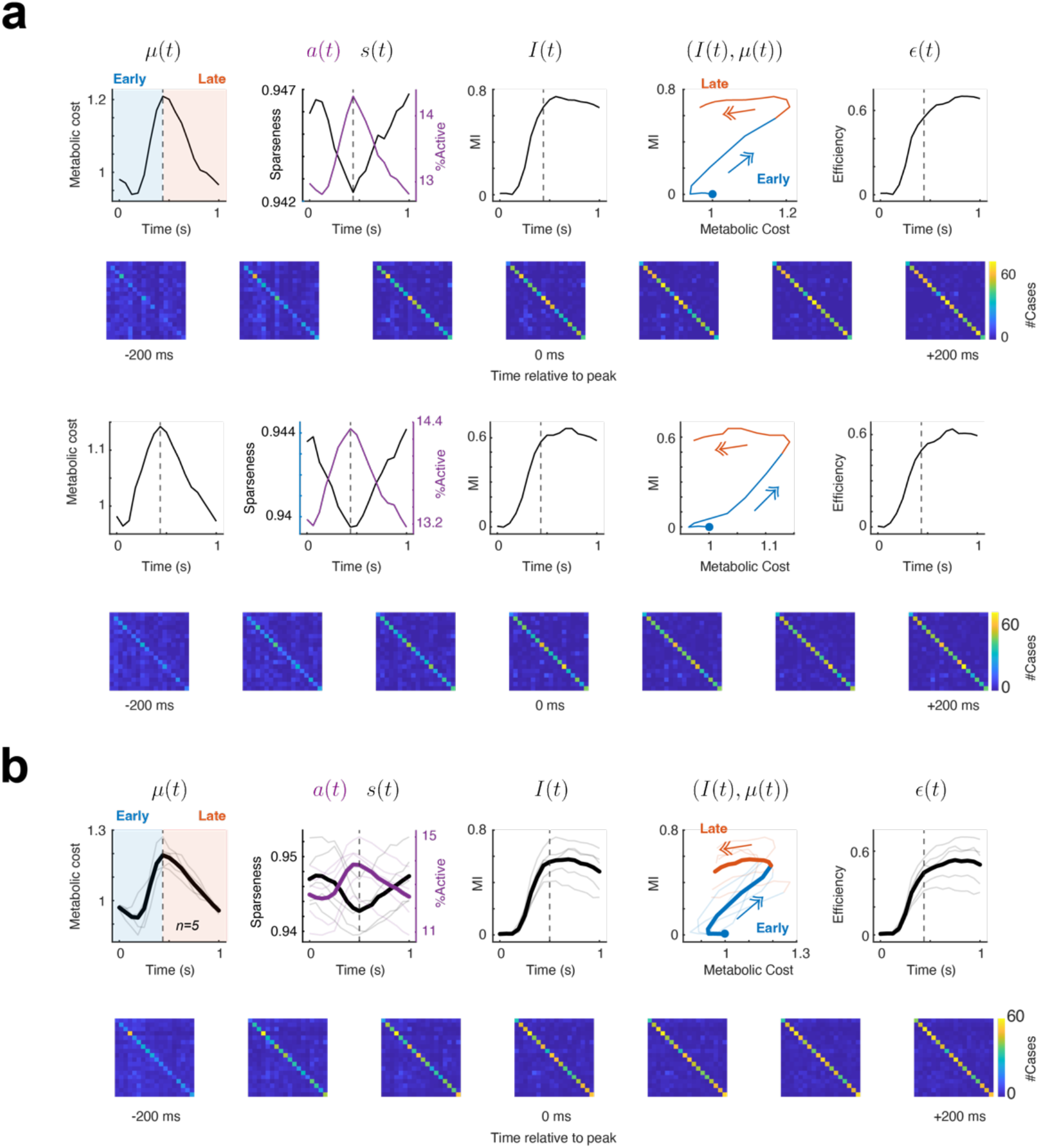
Dynamics of efficient coding using natural movies. **a**. Results in two individual sessions. The format is the same as in **Fig 2. b**. Population data across all the experiments (*n=5* sessions). Transparent curves show the result of individual experiments while the solid curves show the average response across sessions. The images show the average confusion matrices relative to the time to peak.

The dynamic balance between metabolic cost and mutual information can be assessed by plotting the path taken by population activity in the (*μ, I*) − plane during the response period (**Fig 2**). At stimulus onset, the activity starts at (0,1) and rises during the early phase of the response to reach maximum metabolic cost and mutual information at the time to peak (**Fig 2**, blue segment and arrow). In the late phase, the system follows a different trajectory: metabolic cost decreases rapidly, while mutual information declines modestly but remains relatively high (**Fig 2**, red segment and arrow).

The shape of the trajectory in the (*μ, I*) − plane over time demonstrates a key aspect of energy efficiency: there are points in the early and late phases where metabolic cost is identical, while mutual information is significantly higher in the late phase. In other words, the representation becomes more energy-efficient in the late phase compared to the early phase, achieving higher mutual information for the same investment of metabolic cost. Indeed, the dynamics drive the state towards the top left of the graph, locations with high mutual information and low metabolic cost. To quantify this effect, we define the efficiency of the representation as the ratio between mutual information and metabolic cost, *ε*(*t*) = *I*(*t*)/*μ*(*t*). We observe that efficiency increases during the early phase, continues to rise beyond the time-to-peak, and is largely maintained throughout the late phase of the response (**Fig 2**).

A qualitatively similar outcome is observed when the stimuli consist of natural movie sequences **(Fig. 2b**). Metabolic cost shows an initial dip before rising to its peak, while sparseness increases before dipping to a minimum at the same time. The initial dip and the delayed responses compared to grating stimuli likely reflect the lingering effects of the previous stimulus, as the duration of movies was shorter compared to gratings. Still, as with gratings, mutual information rises, peaks, and remains high throughout the response. The trajectory in the (*μ, I*) − plane again reveals that, for the same metabolic cost, mutual information is substantially higher in the late phase. Finally, the efficiency curve for natural movies shows a monotonic increase over time, mirroring the trend observed with synthetic stimuli. The maximum adjusted mutual information is higher for movies than for gratings, likely because movie clips differ substantially from one another, whereas gratings with similar orientations elicit comparable neural responses.

Altogether, our findings reveal that the temporal dynamics of population responses in mouse V1 align with predictions from sparsification networks, providing insights into the adaptive optimization of sensory representations. Following stimulus onset, an initial broad activation is refined over time, leading to increased sparseness, while maintaining mutual information at a high level, thereby improving coding efficiency. We note the response to natural movies is likely influenced by coarse-to-fine spatial frequency dynamics^24,25^, where low spatial frequency signals arrive at the cortex earlier than high spatial frequency information. However, the dynamics of efficiency followed a similar pattern as that observed with gratings that had a fixed spatial frequency.

These results complement earlier studies showing that sparseness increases with the size of the visual stimulation patch^12^ and that population responses to natural images are sparser than responses to synthetic stimuli, enhancing discriminability^15^. While previous work emphasized static sparseness and its dependence on stimulus context, our findings further demonstrate how sparseness, mutual information and coding efficiency evolve dynamically during ongoing sensory processing. These dynamics may serve an evolutionary purpose^26^. In response to novel stimuli, a rapid, high-fidelity response may justify its metabolic cost, particularly when the stimulus could signal a potential threat. Such an energetically costly signal would likely improve evolutionary fitness by facilitating immediate escape responses to aversive stimuli. Once an initial “alarm” is issued, organisms benefit from refining sensory representations to maximize efficiency, conserving metabolic resources while maintaining perceptual fidelity.

One limitation of the study is that two-photon imaging provides limited temporal resolution. We are now using electrophysiological methods to examine the dynamics of these processes at finer temporal scales. Finally, future research should explore the neural mechanisms underlying these dynamics, such as recurrent amplification, lateral inhibition and its spatial extent, or synaptic plasticity, to further elucidate how sparseness and efficiency are regulated in real-time. It will also be of interest to measure population sparseness and efficiency of signals in the LGN to compare to its counterpart in cortical layers 4 and 2+3, to understand the progression of efficient coding along the visual hierarchy.

## Methods

### Experimental model and subject details

All experimental procedures were approved by UCLA’s Office of Animal Research Oversight (the Institutional Animal Care and Use Committee) and were in accord with guidelines set by the U.S. National Institutes of Health. A total of 4 mice, male (3) and female (1), aged P35-56, were used. These animals resulted from cross between TRE-GCaMP6s line G6s2 (Jackson Lab, https://www.jax.org/strain/024742) and CaMKII-tTA (https://www.jax.org/strain/007004). These small numbers did not allow a meaningful comparison between the responses of both sexes.

### Surgery

Imaging was conducted through chronically implanted cranial windows over primary visual cortex. Pre-operatively, mice received Carprofen (5 mg/kg, 0.2 mL after 1:100 dilution) for analgesia. Anesthesia was induced with isoflurane (4%–5%) and maintained during surgery at 1.5%–2%. Core body temperature was kept at 37.5°C, and ophthalmic ointment was applied to protect the eyes. Mice were secured in a stereotaxic apparatus using blunt ear bars positioned in the external auditory meatus. The scalp overlying both hemispheres was removed to expose the skull, which was then dried and coated with a thin layer of Vetbond. After drying (15 minutes), an aluminum bracket was affixed with dental acrylic, and the margins were sealed with Vetbond and additional acrylic to prevent infection.

A craniotomy was performed over monocular V1 on the left hemisphere using a high-speed dental drill, taking care to preserve the integrity of the dura. Once the skull was removed, a sterile 3-mm diameter cover glass was placed directly on the exposed dura and sealed to the surrounding skull with Vetbond. The remaining exposed skull and cover glass margins were further sealed with dental acrylic. Post-surgery, mice were placed on a heating pad until fully awake and then returned to their home cages. Carprofen was administered post-operatively for 72 hours to ensure continued analgesia. Mice were allowed a recovery period of at least six days before the first imaging session.

### Two-photon imaging

Mice were head-restrained on a running wheel and imaged using a resonant two-photon microscope (Neurolabware, Los Angeles, CA) controlled by Scanbox acquisition software and electronics (Scanbox, Los Angeles, CA). Imaging was performed with an Axon 920 laser (Coherent Inc., Santa Clara, CA) at an excitation wavelength of 920 nm. A 16× water-immersion objective (Nikon, 0.8 NA, 3 mm working distance) was used, tilted to align approximately normal to the cortical surface. The microscope operated at a frame rate of 15.6 Hz (512 lines with an 8 kHz resonant mirror), with a field of view measuring 690 μm × 440 μm. Image processing was conducted using a standard pipeline, including image stabilization, cell segmentation, and signal extraction, implemented in Suite2p (https://suite2p.readthedocs.io/).

### Visual stimulation

Visual stimuli were presented on a Samsung CHG90 monitor positioned 30 cm in front of the animal. The screen was calibrated using a Spectrascan PR-655 spectro-radiometer (Jadak, Syracuse, NY), with gamma corrections for the red, green, and blue channels applied via a GeForce RTX 2080 Ti graphics card. Stimuli were generated using a custom-written sketch in Processing 4, leveraging OpenGL shaders (http://processing.org). In the first experiment, sinusoidal gratings with a spatial frequency of 0.04 cycles per degree (cpd) and 90% contrast were presented full-screen, covering 100° × 60° of central vision. Grating orientations were discretized in 10° increments from 0° to 180°, with spatial phases at each orientation uniformly randomized from 0° to 360° in steps of 45°. In the second experiment, visual stimuli consisted of 18 natural movie clips, each 1 second long, sampled from publicly available National Geographic documentaries. These movies were presented full-screen. Stimulus transitions were signaled to the microscope via a TTL line. As an additional failsafe, a small square in the corner of the screen flickered at the onset of each stimulus. The flicker was detected by a photodiode, and its signal was also sampled by the microscope.

### Data Analysis

Denote by 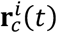 an *N*-dimensional vector 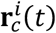 describing the response of a population of *d* neurons to the *i* − *th* trial of a stimulus in class *c* at a time *t* after stimulus onset. Associated with this response, we define 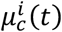 which is the total population activity obtained by adding the entries in 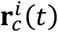. Metabolic cost, *μ*(*t*), is defined as the average of 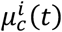 across all stimulus trials and classes. As a measure of sparseness we adopted the Gini index^21^ of the population activity, 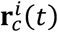, which is denoted by 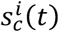. The sparseness signal, *s*(*t*), is then defined as the average of 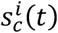 across all stimulus trials and classes. Sparseness ranges from zero (when all the entries have the same value) to one (when all but one entry are zero). Previous studies used 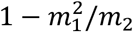 as a measure of sparseness^11,12,19^, where *m*_1_ and *m*_2_ represent the first and second moments of the distribution of activity across the population. We obtain very similar results using this measure (data not shown), so none of our conclusions are impacted by the choice of sparseness measure. We preferred the use of the Gini index because, in our hands, it provides a more robust measure, presumably because it relies on the entire shape of the distribution rather than the first two moments. In addition to the Gini index, we also computed the fraction of active cells, *a*(*t*), defined as the average of 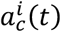 across all stimulus trials and classes. Here, 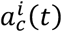 represents the fraction of cells in the population active at time *t*, during the presentation of the *i* − *th* trial of a stimulus in class *c*. We declared a cell “active” if the Suite2p deconvolution signal was larger than zero. Finally, we estimated the mutual information between the stimulus and the response at different times after stimulus onset, *I*(*t*). To compute this quantity, we built a data matrix **X** where each row corresponds to one trial of a stimulus representing the population response *t* sec after the onset of the stimulus. A corresponding column vector **Y** contained the stimulus class for each trial. A *k*-nearest neighbor classifier with leave-one-out cross-validation^27^ armed with a cosine distance metric was used to construct a confusion matrix representing the joint distributions of the true and predicted classes. From the confusion matrix, we can readily estimate the (adjusted) mutual information^22^, *I*(*t*), which ranges from zero (where there is no association between stimulus and response) to one (for perfect association). The optimal value of *k* was determined by looking at the one yielding the maximum mutual information. Note that this method provides a lower bound of the mutual information of the population and the stimulus (due to the data processing inequality).

## Data availability

Raw data describing the response of the population in every single trial along with any ancillary data are available at a Figshare repository at ________________________.

## Code availability

Sample code describing the structure of the database and the replication of our analyses can be found along with the data at ________________________.

## Acknowledgements

We thank Joel Zylberberg and Mario Dipoppa for comments on an earlier version of this manuscript. Supported by EY034488, EY035064, NS116471, EY036219 (DLR).

